# Expression of Fibroblast Growth Factor Receptor 3 (FGFR3) in the Human Peripheral Nervous System: Implications for the Putative Pathogenic Role of FGFR3 Autoantibodies in Neuropathy

**DOI:** 10.1101/2025.03.03.639509

**Authors:** Alexander Chamessian, Diana Tavares-Ferriera, Maria Payne, Raghav Govindarajan, Alan Pestronk, Zach Bertels, Allie J. Widman, Richard A. Slivicki, John S Del Rosario, Jiwon Yi, Bryan Copits, David M. Ornitz, Theodore J. Price, Robert W. Gereau

## Abstract

**Introduction:** Autoantibodies to the Fibroblast Growth Factor Receptor 3 (FGFR3-AAbs) have been associated with idiopathic sensory-predominant neuropathy. The pathogenicity of FGFR3-AAbs in this disorder is unknown. Pathogenic mechanisms of autoantibodies in other dysimmune neuropathies commonly involve their direct binding to antigens on either neurons or glia. The expression of FGFR3 in the human peripheral nervous system is unknown. Therefore, as an initial step toward clarifying the pathogenicity of FGFR3-AAbs, we characterized the expression of FGFR3 in nerve (hNerve), dorsal root ganglia (hDRG) and spinal cord (hSC) in human.

**Methods:** *FGFR3 mRNA* was assayed via in situ hybridization (ISH) on post-mortem sections of hNerve, hDRG and hSC, and by re-analysis of RNA-sequencing data from hDRG. FGFR3 protein was assayed in these tissues using capillary electrophoretic immunoassays (CEIA) with several validated anti-FGFR3 antibodies.

**Results:** *FGFR3* mRNA was not detected in hNerve or hDRG but was abundant in hSC by ISH. FGFR3 protein was absent from hNerve and hDRG by CEIA but was moderately expressed in hSC.

**Discussion:** A direct pathogenic mechanism of FGFR3-AAbs in sensory neuropathy would require the expression of FGFR3 in either neurons or non-neuronal cells in nerve or DRG. Using multiple methods, we did not detect FGFR3 expression at the mRNA or protein levels in these tissues. Given the absence of FGFR3 from hNerve and hDRG, it is improbable that FGFR3-AAbs cause direct damage to the neural components involved in neuropathy and thus are unlikely to be pathogenic, although indirect mechanisms via non-neural cells cannot be excluded.

## Introduction

Autoantibodies against Fibroblast Growth Factor Receptor 3 (FGFR3-AAbs) have been identified in some patients with idiopathic sensory neuronopathy (SNN), small fiber neuropathy (SFN) and sensorimotor polyneuropathy(SMPN)(Nagarajan et al. 2021; J. C. Antoine et al. 2015; Zeidman, Saini, and Mai 2022; Levine et al. 2020). Because of their immmunological origin, FGFR3-AAbs have been proposed to reflect a immune-mediated pathogenesis and on this basis, immunomodulatory interventions such as intravenous immunoglobulin (IVIG), plasma exchange (PLEX) or corticosteroids have been used in some patients with SFN and FGFR3-AAbs, yielding mixed results in the literature. A retrospective case series reported significant improvement in neuropathic pain and intraepidermal nerve fiber density (IENFD)(Zeidman, Saini, and Mai 2022) while a recent double-blind placebo-controlled pilot using IVIG showed neither of these outcomes(Gibbons et al. 2022). Determining the role FGFR3-AAbs would help clarify how to use the presence of these autoantibodies to guide clinical management. The pathogenicity of FGFR3-AAbs has been proposed(Jean-Christophe Antoine 2022) but not demonstrated, neither through passive transfer experiments nor histological analysis.

Pathogenic autoantibodies have been identified in diverse neurological diseases of both the central and peripheral nervous system (PNS)(Prüss 2021) and exhibit multiple mechanisms of action including cross-linking and target receptor internalization, complement activation, stimulatory or inhibitory effects on surface receptors, disrupted protein–protein interaction, antibody-dependent cellular cytotoxicity and disruption of blood–brain or blood-nerve barrier integrity(Prüss 2021). The unifying theme among these diverse mechanisms is the engagement of a specific antigen by the autoantibody within the affected tissue. For FGFR3-AAbs to be pathogenic in sensory neuropathy, it follows that FGFR3 must be expressed in either the peripheral nerve or dorsal root ganglia (DRG). To date there has been no investigation on the presence and distribution of FGFR3 in the human PNS. Therefore, in this study, as an initial step toward addressing the pathogenicity of FGFR3-AAbs, we asked the fundamental question of whether and where mFGFR3 is expressed in the human PNS. Here, we profile the expression of FGFR3 in human DRG, peripheral nerve and spinal cord using multiple complementary methods.

## Methods

### Human Tissue and Serum

Extraction of hDRG, spinal cord and nerve performed as previously described(Valtcheva et al. 2016).

Tissue used for CEIA: Briefly, T11-L5 DRGs, femoral nerves, and lumbosacral spinal cord were extracted from postmortem organ donors in collaboration with Mid-America Transplant. Extraction was performed within 2 hours of aortic cross-clamp. Spinal cord and DRG samples were placed in ice-cold N-methyl-D-glucamine (NMDG)-based solution (93mM NMDG, 2.5mM KCl, 1.25mM NaH2PO4, 30mM NaHCO3, 20mM HEPES, 25mM glucose, 5mM ascorbic acid, 2mM thiourea, 3mM Na+ pyruvate, 10mM MgSO4,0.5mM CaCl2, 12mM N-acetylcysteine; adjusted to pH 7.3 using NMDG or HCl, and 300-310 mOsm using H2O or sucrose) for transport. Femoral nerves were stored without buffer in a clean container with moistened gauze. Extracted tissues were cleaned and then snap frozen on dry ice and stored at -80ºC for long-term storage.

Tissue used for RNAscope: Human lumbar DRG and spinal cord tissues were collected from organ donors within 4 hours of cross-clamp and from neurologic determination of death donors in collaboration with Southwest Transplant Alliance in Dallas, TX. Tibial nerve sample was recovered from diabetic patients undergoing amputation surgery. Samples were randomly selected on the basis of sex and age from the tissue repository at University of Texas at Dallas. All human tissue procurement procedures were approved by the Institutional Review Boards at the University of Texas at Dallas.

Serum: Serum containing FGFR3-AAbs from a female patient with biopsy-confirmed small fiber neuropathy was obtained from the Neuromuscular Clinical Laboratory at Washington University School of Medicine in St. Louis. FGFR3-AAbs were detected using previously described methods (Levine et al. 2020).

### Animals

C57BL6/J male mice aged 8-12 weeks were used for RNAscope. Briefly, animals were transcardially perfused with phosphate-buffered saline (PBS) followed by 4% paraformaldehyde-PBS. Tissues were extracted including DRG and lumbosacral spinal cord. The tissues were post-fixed for 2 hours and then cryoprotected in 30% sucrose-PBS solution at 4<u>º</u>C until the tissues were fully saturated.

### Capillary Electrophoresis Immunoassay (CEIA)

Protein extracts from human DRG, nerve and spinal cord were prepared using the Precellys 24 (Bertin Instruments) and CK-14 ceramic beads. RIPA buffer (Millipore-Sigma) supplemented with protease inhibitors (Roche; #11836170001) and phosphatase inhibitors (Halt Protease Cocktail, ThermoFisher Scientific; #78420) was combined with tissue (∼100 mg) in the CK-14 tubes and homogenization was performed using a program of 5000 rpm x 2 cycles x 20s with 30s delay. The tissue protein lysates were clarified by centrifugation and then stored at -80<u>º</u>C until further use. Protein lysates or recombinant human FGFR3 (ThermoFisher Scientific, #PV3145) were diluted in 0.1x Simple Western sample buffer (Protein Simple) and then denatured at 70<u>º</u>C for 5 mins. Primary and secondary Antibodies were used at the following concentration: Rabbit anti-FGFR3 (1:50, Cell Signaling Technologies, RRID:AB_2246903), Rabbit anti-FGFR(1:50, Abcam, RRID:AB_2810262), Rabbit anti-FGFR3(1:50, Santa Cruz Biotechnology, RRID:AB_627596), anti Anti-Rabbit-HRP (undiluted, Protein Simple, RRID:AB_2860577). Simple Western CEIA was performed on the Wes instrument (ProteinSimple) according to the manufacturer’s instructions. Data were analyzed using the Compass (v.6.1) Software (ProteinSimple).

### In Situ Hybridization (ISH)

Human tissues: hDRG, nerve and spinal cord from UT Dallas was mounted in Tissue-Tek O.C.T embedding medium (Sakura), cryosectioned and thaw-mounted on Superfrost slides (VWR). RNAscope 2.5 HD Assay Red (ACD Bio) was performed according to the manufacturer’s instructions using probes Hs-FGFR3(#310791), Hs-PRPH(#410231), and Negative Control Probe-DapB(#310043). The slides were counterstained with hematoxylin after ISH. All slides were imaged on a widefield microscope (Olympus).

Mouse tissues: Cryoprotected mouse DRG and spinal cord were embedded in Tissue-Tek O.C.T and cryosections were made at 12*μm* and 14 *μm*, respectively, and thaw-mounted on Superfrost slides. RNAscope Multiplex Fluorescent V2 assay was performed according to the manufacturer’s instructions using the ‘Fresh Frozen’ pretreatment protocol consisting of incubation with protease IV for 30 min at room temperature. The following probes were used: Mm-Fgfr3 (#440771), Mm-Nefh-C3 (#443671-C3). After ISH, slides were counterstained with 4’,6-diamidino-2-phenylindole (DAPI) and mounted with. Following RNAscope, slides were imaged using Leica DM6b system at 20x magnification with the Leica Application Suite X (LAS-X, v.3.7, Leica Microsystems). Acquisition setting were kept consistent throughout imaging. Images were analyzed in Fiji.

### Transcriptome Analysis

RNA-seq data were sourced from Ray et al.(Ray et al. 2018), Wangzhou et al.(Wangzhou et al. 2020), and North et al.(North et al. 2019) and represented as transcripts per million (TPM) to enable comparison between different datasets and tissues. Full details of RNA-seq data processing can be found in their originating publications. Analysis of the data was performed using R(Team 2023) with the following packages: ggplot2(Wickham 2016).

## Results

### Expression of *FGFR3* mRNA in human dorsal root ganglia, nerve and spinal cord

To assess the expression of *FGFR3* mRNA in tissue, we conducted chromogenic *in situ* hybridization (RNAscope) on human lumbar dorsal root ganglia (DRG),nerve (femoral) and lumbosacral spinal cord (SC). We observed no *FGFR3* expression in DRG or nerve, and obtained expected results using the negative control probe (Fig. 1A-D). In contrast, *FGFR3* was clearly expressed in the central gray matter of SC(Fig 1E-G). Although we did not perform double-staining, the *FGFR3* signal observed in the spinal cord is most likely localized to astrocytes, based on recent single-cell transcriptomics findings in the CNS demonstrating strong *FGFR3* expression in astrocytes(Hodge et al. 2019; Yadav et al. 2023). We also performed single-molecule fluorescence *in situ* hybridization in mouse tissues and similarly found no expression of *Fgfr3* mRNA in DRG (mDRG) but abundant expression in the gray matter of the spinal cord (mSC) (Fig. 2).

**Figure 1.**
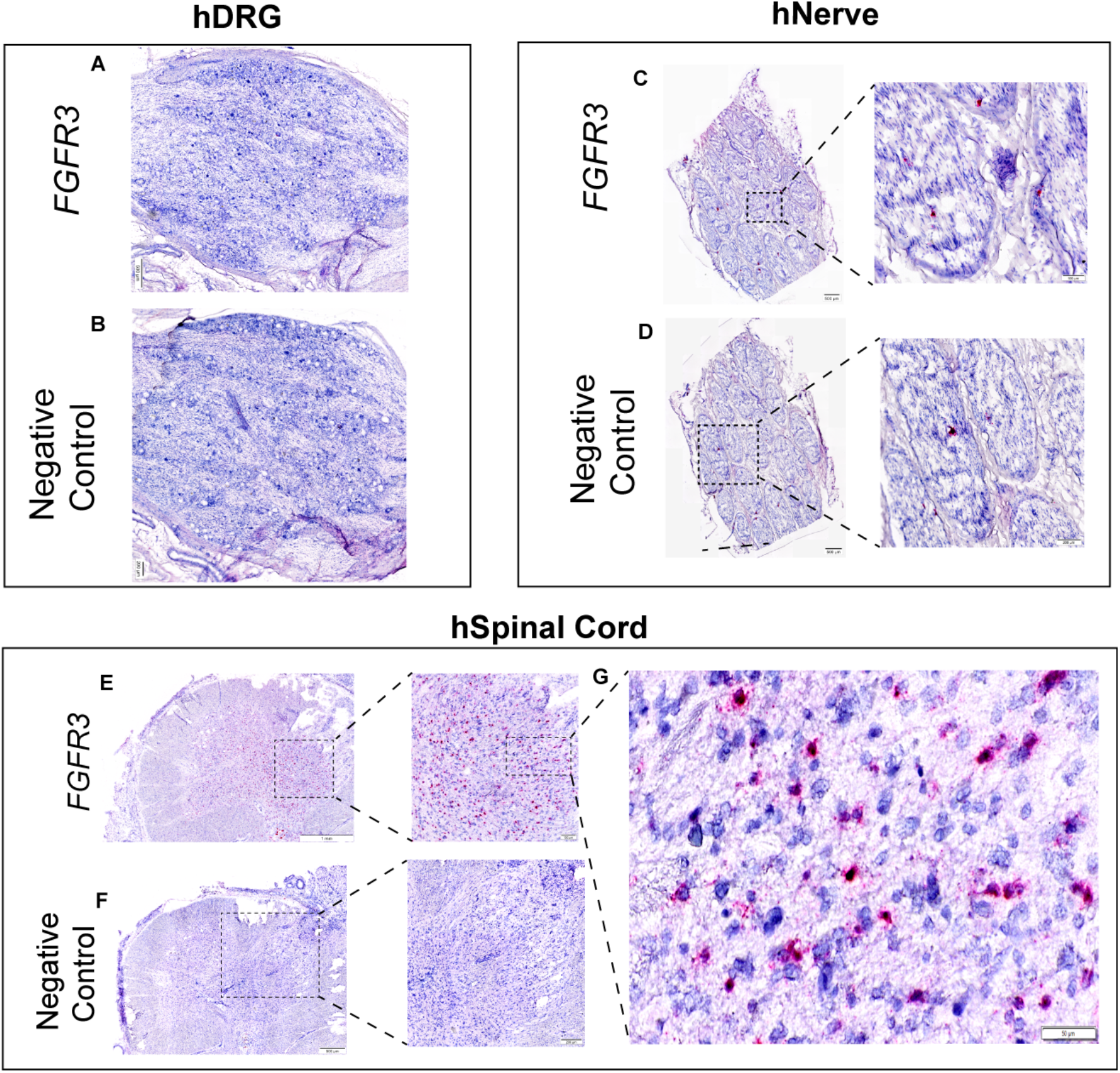
Distribution of FGFR3 mRNA in human neural tissues. Representative images of chromogenic in situ hybridization (RNAscope) of FGFR3 in DRG (**A**) and negative control probe (**B**). Representative images of FGFR3 RNAscope in tibial nerve (**C**) and negative control probe (**D**). Representative images of FGFR3 RNAscope in human lumbosacral spinal cord (**E**) and inset (**G**). Negative control probe in human spinal cord (**F**).

**Figure 2.**
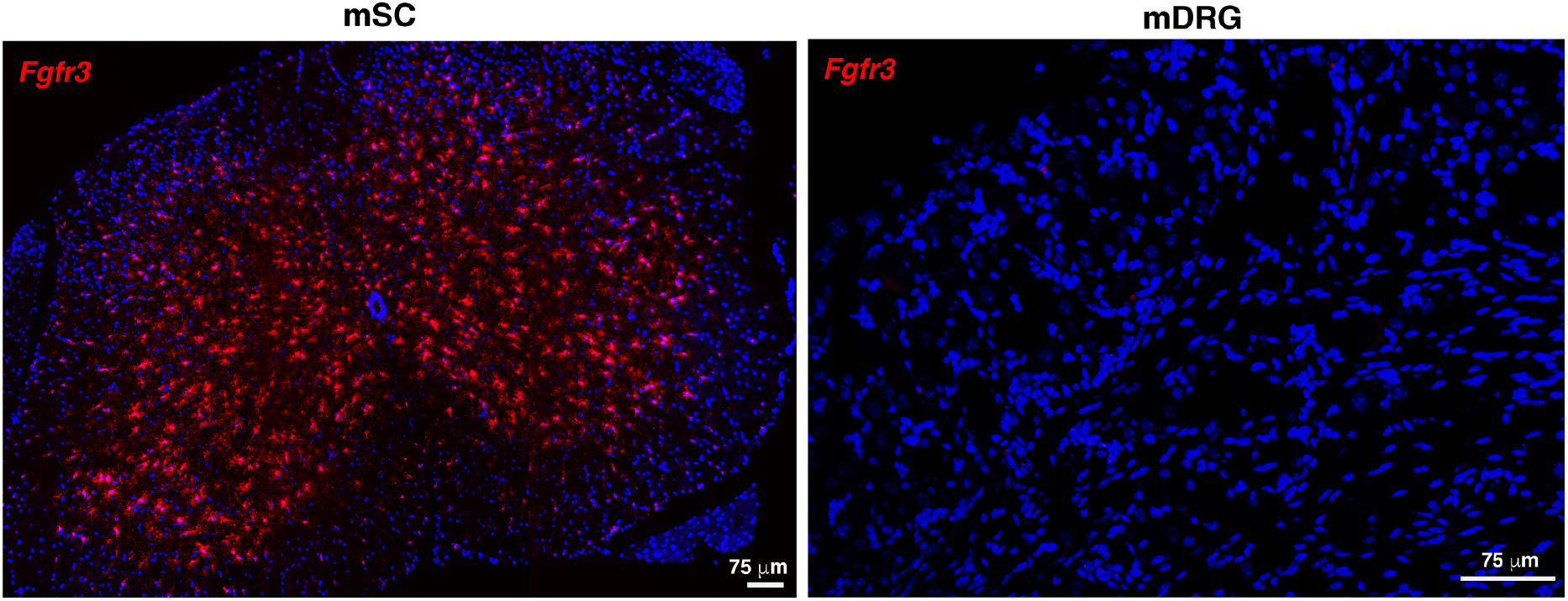
Distribution of Fgfr3 mRNA in mouse spinal cord (mSC, left) and dorsal root ganglion (mDRG, right). Red = Fgfr3; Blue = DAPI.

To further extend these observations, we analyzed RNA-seq datasets from our previous studies of human nervous system tissues(Ray et al. 2018) as well as publicly available data (GTEx)(Lonsdale et al. 2013) to assess FGF receptor (FGFR) expression. *FGFR3* is nearly undetectable in DRG, with an average transcript per million (TPM) value of 0.7, consistent with a recent study of spatial sequencing of human DRG where *FGFR3* was barely detectable(Tavares-Ferreira et al. 2022). In tibial nerve, the average TPM is 4.8, while in SC, *FGFR3* has an average expression of 23.3 TPM, consistent with our RNAscope observations (Fig. 3A). Notably, frontal cortex has an average expression of 189.9 TPM, reflecting its known expression in astrocytes(Hodge et al. 2019).

**Figure 3.**
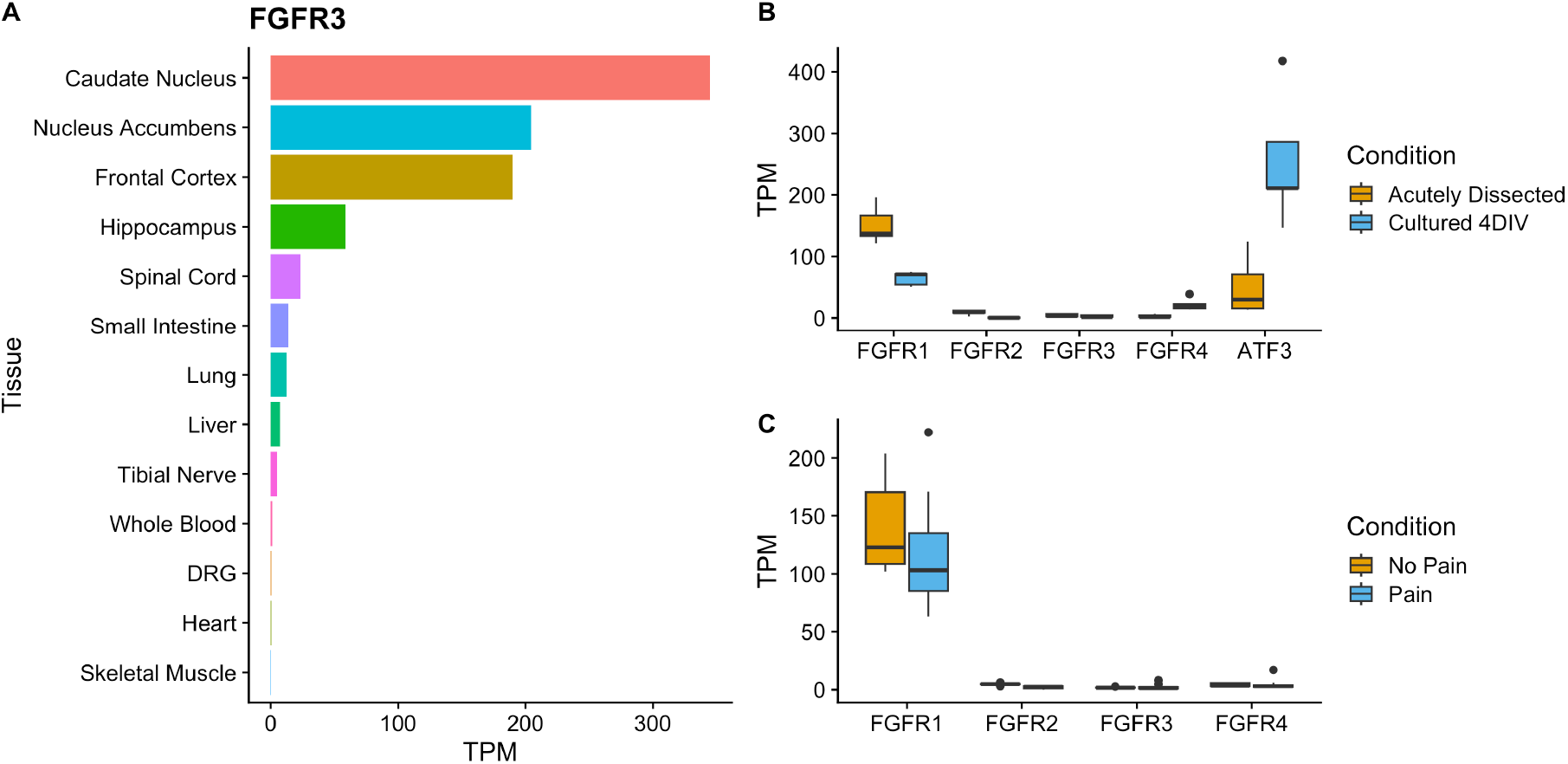
RNA-seq of FGFRs in human tissues. (A) Transcript abundance of FGFR3 expressed in transcripts per million (TPM) in selected human tissues. (B) FGFR family members and ATF3 transcript abundance in human human dorsal root ganglia (hDRG) primary culture. (C) FGFR family member transcript abundance from whole hDRG tissues from patients with and without pain. Data for (A) from (Ray et al. 2018), (B) (Wangzhou et al. 2020), and (C) (North et al. 2019).

Having established that *FGFR3* is nearly absent from normal hDRG, we asked whether *FGFR3* expression could be induced by external insults, given that previous reports in mouse suggested that nerve injury upregulates *Fgfr3* in mDRG(Jungnickel et al. 2005). To that end, we re-analyzed previous RNA-seq data of dissociated cultures of human DRG (Wangzhou et al. 2020) as well as whole DRG from patients with and without pain who underwent resection of a DRG in the setting of surgery for spinal malignancy(North et al. 2019). In hDRG culture, *FGFR3* average expression immediately after dissociation was 4.1 ± 1 TPM, and 2.8 ± 0.7 TPM on day *in vitro* (DIV) 4, showing no upregulation in response to dissociation (Fig. 3B). Consistent with the fact that dissociation represents an injured state, the transcription factor *ATF3*, which is known to increase dramatically in response to nerve injury(Cheng et al. 2021), was upregulated more than five-fold at DIV4, with an average expression value of 254.6 ± 46.4 TPM compared to 50.9 ± 21.1 TPM at baseline (Fig. 3B). Similarly, in surgically-resected DRGs, *FGFR3* remained virtually undetectable between painless and painful groups (Fig. 3C).

### FGFR3 protein expression human dorsal root ganglia, nerve and spinal cord

To assess the protein-level expression of FGFR3, we conducted a high-sensitivity capillary electrophoresis immunoassay (CEIA, Simple Western) on human lumbar dorsal root ganglia (DRF), nerve (femoral) and lumbosacral spinal cord (SC) protein extracts. Given our findings of abundant *FGFR3* mRNA expression in the spinal cord, we used hSC protein extracts to evaluate several commercially available anti-FGFR3 antibodies and identified a rabbit monoclonal antibody (FGFR3-mAB, Cell Signaling Technologies) targeting the intracellular domain of FGFR3 that demonstrated excellent performance with a single peak at the expected molecular weight (Fig. 4A). Furthermore, we confirmed the specificity and evaluated the sensitivity of the FGFR3-mAb by assaying purified recombinant human FGFR3 intracellular (rFGFR3-IC) domain, which is the same antigen used in the clinical detection of FGFR3 autoantibodies from patients with neuropathy using ELISA(J. C. Antoine et al. 2015). In this way, we detected clear and strong binding of the FGFR3-mAb antibody to purified FGFR3 intracellular domain with an input range from 160 pg down to 0.8pg (Fig. 4B). We also probed the purified rFGFR3-IC with serum from a patient with sensory neuropathy and FGFR3-AAbs as detected by ELISA (Fig. 4C). Using a 1:20 dilution of serum, we observed a single peak at 57 kDa corresponding to the rFGFR3-IC, demonstrating that patient-derived FGFR3-AAbs and the FGFR3-mAb both recognize the purified rFGFR3-IC domain in the capillary immunoassay. Finally, having validated the FGFR3-mAB in multiple ways, we assayed protein extracts from three separate donors in the CEIA and observed no FGFR3 protein expression in hDRG or hNerve but strong expression of FGFR3 in hSC (Fig. 4D).

**Figure 4.**
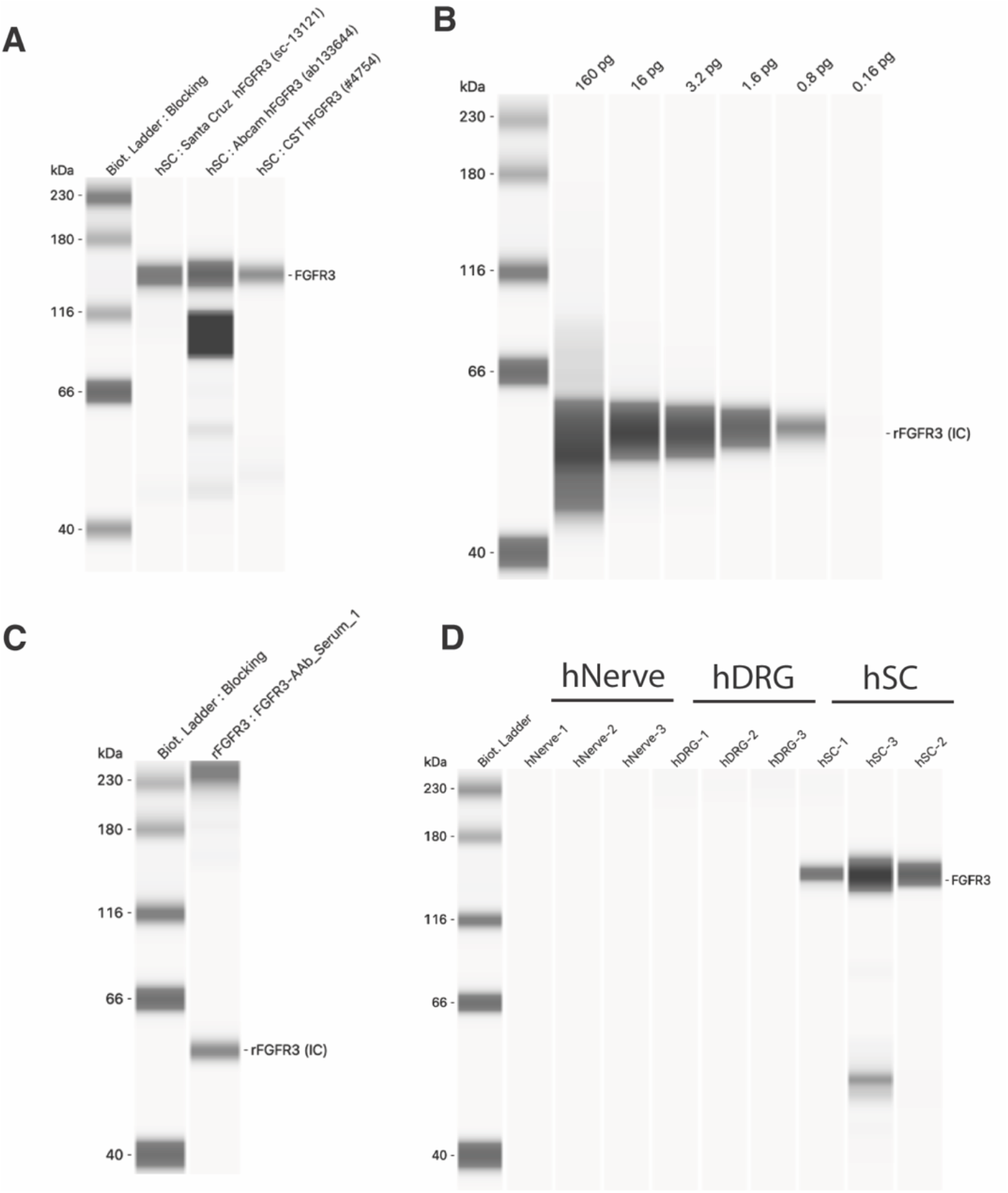
Capillary Electrophoresis Immunoassay (CEIA) of FGFR3 from human tissues. (A) Evaluation of anti-FGFR3 commercial antibodies in CEIA of hSC protein extracts. (B) Validation of FGFR3-mAB against purified recombinant FGFR3 intracellular domain (rFGFR3-IC) in CEIA at varying input amounts (pg: picogram). (C) CEIA of rFGFR3-IC using patient-derived FGFR3-AAbs. (D) CEIA showing FGFR3 abundance in protein extracts from human femoral nerve (hNerve), hDRG and hSC, n=3 donors per tissue.

## Discussion

The studies performed here demonstrate that FGFR3 is not expressed in nerve and dorsal root ganglia in humans in both normal and pathological conditions. Using *in situ* hybridization, we demonstrated that *FGFR3* mRNA was virtually undetectable in nerve and DRG, which was corroborated by re-analysis of several RNA-seq datasets from our previous work. Consistently, we also show that FGFR3 protein is undetectable in DRG and nerve by targeted immunoassay. Importantly, we did observe *FGFR3* mRNA and FGFR3 protein in SC, demonstrating the ability of the multiple assays we used to detect FGFR3 if it is present. Among the four FGFRs, FGFR1 was by far the most abundant at the mRNA levels in DRG and nerve, and FGF1 and FGF2 were the most abundant ligands, suggesting that any FGFR signaling in hDRG and nerve occurs through these receptor-ligand pairs, at least under non-pathological conditions. Our findings contrast with those of Antoine *et al*. where immunohistochemistry of adult rat dorsal root ganglia and embryonic rat trigeminal placode were used to substantiate the expression of FGFR3 in sensory neurons. In light of the multiple lines of evidence we present here showing no little to no expression of FGFR3 in the mouse and human DRG, it is likely that the immunostaining in Antoine et al. was non-specific.

The pathogenicity of FGFR3-AAbs has been questioned on the grounds that its epitope is contained in the intracellular domain of FGFR3. However, this fact does not preclude the potential for FGFR3-AAbs to be pathogenic, given that other autoantibodies with intracellular targets such as anti-Hu have been shown to cause neuronal death(Greenlee et al. 2014). An even more fundamental question is whether the antigen of an autoantibody is present at all in the involved tissue or structure. Accordingly, the principal motivation of our study was to characterize the expression of FGFR3 in the human peripheral nervous system in order to ultimately address the question of whether and how FGFR3-AAbs contribute to sensory neuropathy.

Given our central finding of absence of FGFR3 protein from hNerve and hDRG, it is improbable that FGFR3-AAbs are directly pathogenic in the setting of sensory neuronopathy and/or neuropathy. Pathogenic autoantibodies have been demonstrated in numerous neuropathies such as Guillain Barre syndrome (GBS), chronic inflammatory demyelinating polyneuropathy (CIDP), anti-MAG neuropathy, and Multi-Focal Motor Neuropathy (MMN)(Martín-Aguilar, Pascual-Goñi, and Querol 2019; Pascual-Goñi et al. 2021; Dalakas 2015). In all cases, the pathogenesis involves binding of the autoantibodies to antigens present on neurons, their axons, ensheathing Schwann cells. To our knowledge, there have been no demonstrations of an autoantibody playing a pathogenic role in neuropathy where the antigen is not present in the target tissue.

The nerve and DRG samples that we analyzed in this study are from donors without neuropathy, so these would be considered normal, non-pathological specimens. Could some pathological stimulus induce the expression of FGFR3 in DRG or nerve such that FGFR3-AAbs could engage their antigen? We cannot exclude this possibility, but based on our RNA-seq analysis of cultured and whole hDRG, this is unlikely. Cultured DRG cells represent an injured state given that the dissociation process axotomizes sensory neurons, as reflected by the marked increase of *ATF3*. Under these conditions, *FGFR3* expression did not change compared to baseline. Similarly, whole DRG from patients with spinal malignancy, where the DRG is in close proximity to the malignant process or in some cases directly compressed by it, there also was no increase in *FGFR3* expression. Finally, in studies in mice, nerve injury, chemotherapy treatment and inflammation did not induce *Fgfr3* expression, as it was nearly undetectable in all experiments in the mDRG across models(Renthal et al. 2020). It may still be the case that some other stimulus besides these could induce the expression of FGFR3 in DRG or nerve but it is unclear what that might be and how it would differ so markedly from those that we evaluated. Indirect mechanisms involving non-neural cells in target tissues (e.g. skin), infiltration of non-resident immune cells into DRG or nerve, or long-range systemic neuro-immune interactions are conceivable albeit highly speculative and improbable.

What then to make of the role of FGFR3-AAbs in neuropathy? Our findings argue against a pathogenic role for these autoantibodies and lead to the conclusion that FGFR3-AAbs are likely an epiphenomenon, as others have suggested(Kovvuru et al. 2020). Nevertheless, even if they are not pathogenic, FGFR3-AAbs hold demonstrated value as diagnostic biomarkersMoritz et al. (2020) and may still reflect a dysimmune pathogenesis involving other immune effector mechanisms. Future investigations will be needed to elucidate the causes and mechanisms of neuropathy associated with FGFRS-AAbs. When such studies are undertaken, our results here should help to guide experimental design. For example, given the demonstrated absence of FGFR3 protein from DRG and nerve, any *in vivo* or *in vitro* outcomes arising from the use IgG fraction from patients with FGFR3-AAbs (as is standard in such work) should be interpreted with caution. Ideal experimental design would employ affinity-purified FGFR3-AAbs to ensure the specificity of any observed effects.

Our study has several limitations. We include only three human tissue donors in our protein analysis, and one donor for hDRG/hSC and hNerve for *ISH* analysis. Furthermore, we were unable obtain DRG or nerve tissue from patients with FGFR3 antibody-related neuropathy, preventing us from testing directly whether there is FGFR3 expression in these tissues in the setting of disease.

In conclusion, our study demonstrates that FGFR3 is very lowly expressed in the human PNS under normal and pathological conditions, and consequently, the pathogenicity of FGFR3-AAbs in neuropathy is improbable.

## 0.1 Acknowledgements

We would like to thank the organ donors and their families for their gift that has enabled this research. We thank Lawrence A. Zeidman for his helpful feedback on this manuscript.

## 0.2 Author Contributions

A.C.: Conceptualization, Investigation, Formal Analysis, Funding Acquisition, Visualization, Writing - original draft, Supervision D.T.: Investigation, Visualization, Formal Analysis, Writing - review & editing T.J.P.: Funding, Writing - review & editing, Supervision. M.P.: Investigation R.G.: Conceptualization, Writing - review & editing A.P.: Resources Z.B.: Resources A.W.: Resources R.A.S: Resources J.S.D: Resources J.Y.: Resources D.M.O.: Conceptualization, Funding Acquisition, Writing - review & editing

## 0.3 Funding

Funding for this work was provided by Hope Center for Neurological Disorders Pilot Grant (RG), U19NS130608 (TJP)

## 0.4 Declarations

The authors declare that they have no conflict of interest

## Notes

**Conflict of Interest Statement:** The authors have no conflict of interests to declare.

### Competing Interest Statement

The authors have declared no competing interest.

